# Transcriptome variations in hybrids of wild emmer wheat (*Triticum turgidum ssp. dicoccoides*)

**DOI:** 10.1101/2023.10.23.563532

**Authors:** Alon Ziv, Khalil kashkush

## Abstract

**Background:** Wild emmer wheat is a great candidate to revitalize domesticated wheat genetic diversity. Recent years have seen intensive investigation into the evolution and domestication of wild emmer wheat, including whole-genome DNA and transcriptome sequencing. However, the impact of intraspecific hybridization on the transcriptome of wild emmer wheat has been poorly studied. In this study, we assessed changes in methylation patterns and transcriptomic variations in two accessions of wild emmer wheat collected from two marginal populations, Mt. Hermon and Mt. Amasa, and in their stable F4 hybrid.

**Results:** Methylation-Sensitive Amplified Polymorphism (MSAP) detected significant cytosine demethylation in F4 hybrids vs. parental lines, suggesting potential transcriptome variation. After a detailed analysis, we examined nine RNA-Seq samples, which included three biological replicates from the F4 hybrid and its parental lines. RNA-Seq databases contained approximately 200 million reads, with each library consisting of 15 to 25 million reads. There are a total of 62,490 well-annotated genes in these databases, with 6,602 genes showing differential expression between F4 hybrid and parental lines Mt. Hermon and Mt. Amasa. The differentially expressed genes were classified into four main categories based on their expression patterns. Gene ontology (GO) analysis revealed that differentially expressed genes are associated with DNA/RNA metabolism, photosynthesis, defense response, and phosphorylation.

**Conclusion:** This study highlights the significant transcriptomic changes resulting from intraspecific hybridization within natural plant populations, which might aid the nascent hybrid in adapting to various environmental conditions.

## Background

Wheat is one of the most important crops for human consumption, playing a significant role in global food production [1]. Ensuring a higher yield of wheat is crucial to maintaining global food security [2]. Global wheat production in 2021 reached 777 million tons according to the Food and Agriculture Organization of the United Nations (http://www.fao.org/faostat/en/#home). Modern agriculture prioritizes pure breeding, leading to a narrow gene pool among wheat species despite high yield [3–5]. The loss of genetic diversity in wheat has increased vulnerability to diseases, pests, and climate change, posing a threat to communities and countries [4–7]. Without adaptation to climate change, crop production is predicted to decrease with a 2°C temperature increase from late 20th-century levels, according to the IPCC [8]. During certain wheat growth stages, drought can cause up to a 92% loss in yield [9]. Moreover, certain diseases like Ug99, a fungal infection that caused severe damage to wheat cultivation in Africa and the Middle East, pose a significant danger to global food security, and adversely affect wheat growth, leading to a massive impact on food security and human health all around the world [10].

A potential source to revive genetic diversity in domesticated wheat is wild emmer wheat, the progenitor of modern wheat. The genetic diversity of wild emmer wheat has been preserved, allowing it to remain adaptive to various environmental and biotic stressors [3, 11, 12]. The high genetic diversity of wild emmer wheat makes it a crucial subject for crop research, as it provides highly sought-after traits and mechanisms for environmental adaptation and stress resistance. In Israel, the wild emmer’s habitat spans from Mt. Hermon to Mt. Amasa in the Judean Desert [13, 14]. Although it is geographically small, this region boasts a diverse range of habitats, each with its unique ecological conditions. These habitats vary in elevation, ranging from 200 meters below sea level in the Jordan Valley to 1600 meters above sea level at Mt. Hermon.

Additionally, they differ in soil types, temperatures, and other biotic and abiotic conditions [13, 15]. Previous studies have primarily focused on the genetic diversity of wild emmer wheat populations within Israel, at both macro and micro scales, with findings suggesting correlations between genetic diversity, ecological traits, and geographical location [11, 12, 14, 16–18]. The distribution of wild emmer wheat populations in Israel was influenced by population position (core or marginal) and size [14]. Additionally, studies on Israeli emmer populations have revealed non-random genetic differentiation linked to soil, topography, and climate at both single and multi-locus levels [11]. Finally, the study by Venetsky et al., (2015) detected population-specific methylation patterns in the whole genome and around TEs in five geographically isolated populations in Israel. Furthermore, these population-specific methylation patterns were heritable and passed down from the first to the second generation. Together with the phenotypic plasticity of wild emmer wheat and its population spread across a wide range of habitats in Israel, these findings suggest underlying adaptive genetic and epigenetic mechanisms.

Whole-genome sequencing breakthroughs have greatly improved the sequencing of wheat genomes. The draft sequence of wild emmer wheat [19] provided an opportunity to investigate and assess transcriptomic changes within natural populations, revealing functional adaptations to different environments. For example, Yin et al., [20] identified and isolated a single dominant powdery mildew resistance gene from the wild emmer wheat natural population found at Mt. Carmel in Israel, showing the possibilities and opportunities for domesticated wheat improvement can be found in the wild emmer wheat.

In this study, we aimed to explore changes in methylation patterns and transcriptomic variations in wild emmer wheat accessions and their hybrid. We selected parental lines from marginal Israeli wild emmer wheat populations - Mt. Hermon and Mt. Amasa - and their stabilized F4 hybrid. To determine the changes in epigenomic patterns, we utilized MSAP (Methylation-Sensitive Amplified Polymorphism), which is a modified version of the typical AFLP, to estimate methylation patterns on a genome-wide scale. For analyzing transcriptomic variations, we conducted RNA-Seq analysis. We then characterized the differentially expressed genes between the parental lines and their F4 hybrid at the molecular level. Additionally, we analyzed and discussed the functions and biological pathways of the differentially expressed genes.

## Results

### Cytosine methylation status in parental lines and their hybrid

Cytosine methylation patterns were examined to determine whether significant epigenetic alterations exist between hybrids and parental lines. We utilized MSAP to determine the cytosine methylation status at CCGG sites. Our study involved the analysis of twelve plants to compare the methylation differences between parental lines and F4 hybrid. For each of the parental lines, Mt. Hermon and Mt. Amasa, we conducted three biological replicates, while for the F4 hybrid, we conducted four replicates. To this end, we conducted three MSAP reactions using different selective primer combinations of EcoRI/MspI and EcoRI/HpaII (primer sequences are listed in Table S1, Additional file 1).

In each MSAP reaction, we analyzed 200-300 peaks comprising 740 CCGG sites (265 bands in E-ACA/HM-TCAA, 200 bands in E-ACT/HM-TCAA, and 275 bands in E-ACC/HM-TCAA). Methylation levels were calculated by dividing the number of polymorphic bands between treatments (*HpaII* and *MspI*) by the total band count. The presence of a monomorphic MSAP site in both *MspI* and *HpaII* patterns indicates an unmethylated CCGG site. Polymorphic bands are observed when a band appears in the *MspI* pattern, but not in the *HpaII* pattern. This indicates that the internal cytosine in that particular CCGG site is methylated. Conversely, when a band appears in the *HpaII* pattern, but not in the *MspI* pattern, it indicates that the external cytosine in the same CCGG site is methylated in one of the DNA strands. This is known as hemi-methylation status. Note that methylation of both cytosines at the CCGG site prevents PCR amplification in both *MspI* and *HpaII* patterns. The Hermon parental line had an average cytosine methylation level of ∼85.6%, while the Amasa parental line had an average methylation level of ∼82.4% (see Table S2, Additional file 1). The methylation level in F4 hybrids was approximately 75%, suggesting demethylation of CCGG sites in hybrids compared to parents. This may be associated with higher genome expression.

Alterations in DNA methylation patterns at specific CCGG sites were identified by analyzing the difference in band composition between parental lines and hybrids. To illustrate, a polymorphic band that was present only in the parental lines and not in F4 hybrids was considered a demethylation event in the F4 hybrids. Conversely, a polymorphic band found only in F4 hybrids indicated a hypermethylation event (see supplemental Fig. S1, Additional file 2 for an example of such alteration). Detailed analysis showed demethylation of 50 CCGG sites and hypermethylation of 26 sites in the F4 hybrid, indicating a significant decrease in methylation.

### Transcriptome analysis in the parental lines and their F4 hybrid

To evaluate transcriptome changes in wild emmer wheat hybrids and their parental lines, we performed RNA-Seq analysis on an F4 hybrid and its parents.

#### Library quality

Leaf total RNA samples collected from the parental lines, Mt. Hermon (MH) and Mt. Amasa (MA), as well as their F4 hybrid, were fully sequenced. To obtain comprehensive coverage and perform accurate statistical analysis, we sequenced three biological replicates of the leaves simultaneously. As part of this effort, we collected RNA-Seq data from 9 libraries (3 from each of the two parents and 3 from the F4 hybrid), with each library containing between 15 to 25 million reads, resulting in a total of approximately 200 million reads. Alignment of the RNA-Seq data to the reference RNA-seq data of WEWSeq 1.0 acc. Zavitan resulted in a mapping rate of approximately 82% for each of the three libraries. A total of 62,490 genes were identified in the parental and hybrid libraries, with ∼38,000 genes in common.

#### Variations in gene expression patterns

The study used Principal Component Analysis (PCA) to compare the gene expression patterns between the parental lines and their F4 hybrid. The analysis was conducted on normalized gene counts of more than 38,000 genes identified through RNA-Seq. The results showed that the first and second components (PC1 and PC2) explained 63.38% and 21.53% of the total variation in the transcriptome (Fig. 1). PCA analysis separates the parental lines and F4 hybrid (Fig. 1). F4 hybrid was positioned between the parental lines on the PC1 axis, but appeared less similar to them on the PC2 axis (Fig. 1). It was observed that the main differences within the groups were found in F4 and Mt. Amasa. On the PC1 and PC2 axes, Mt. Amasa showed the most significant variation within the group, which was more pronounced on the PC1 axis. On the other hand, F4 exhibited relatively high variation within the group over the PC2 axis. This within-group variation was also noticed when differentially. In addition, the heatmap (Fig. 2) displaying the total transcriptome expression value per sample supported the PCA analysis. The heatmap suggests that the F4 hybrids are positioned between the parental lines and are slightly closer to Mt. Hermon. Moreover, it highlights the within-group variation observed in both F4 and Mt. Amasa to a greater extent.

**Fig. 1.**
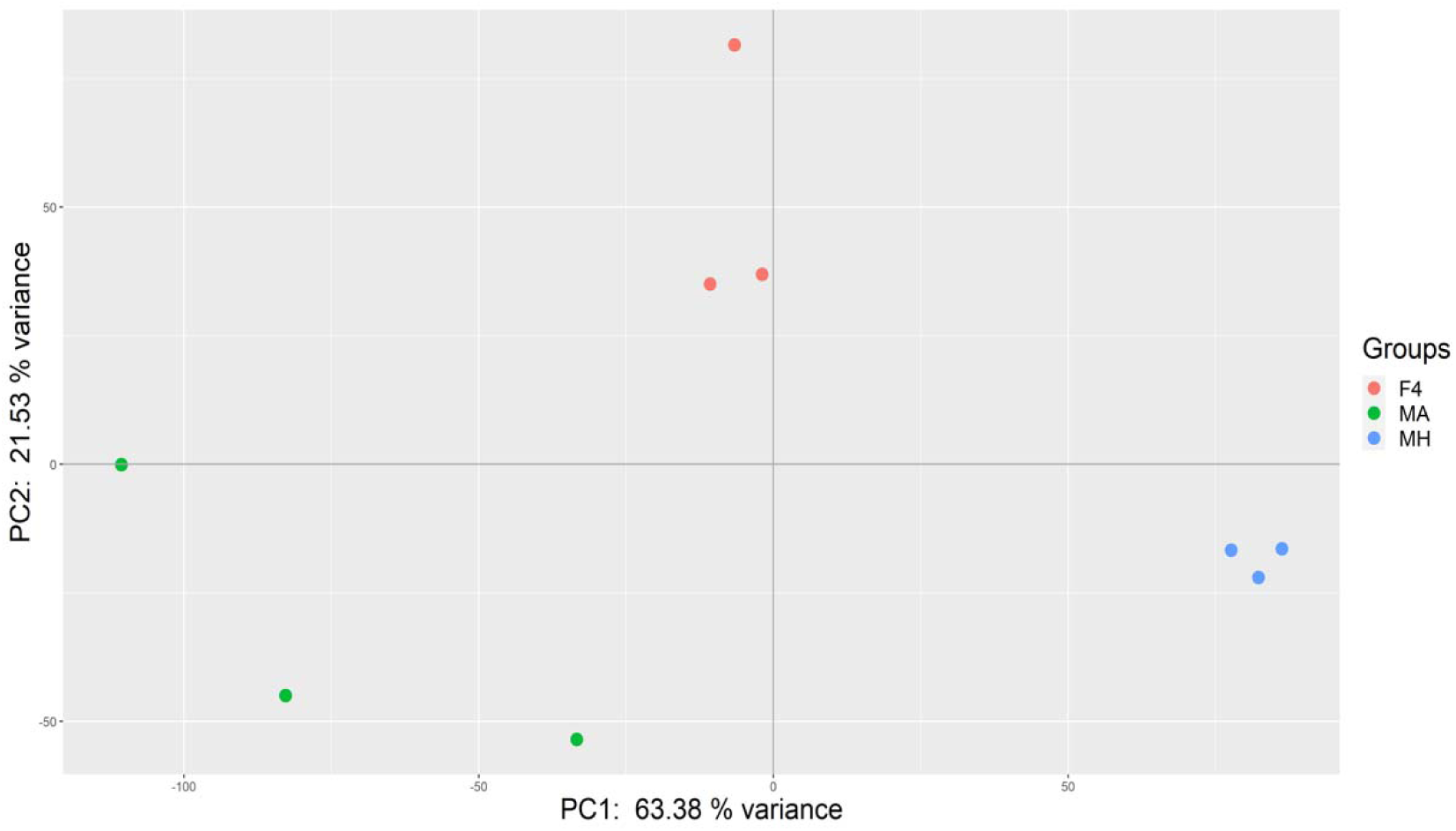
PCA plot of F4 hybrids and Mt. Amasa (MA) and Mt. Hermon (MH) parental lines. PCA was generated from over 38,000 genes identified through RNA-Seq. PC1 explains 64% of the variance, separates between the groups F4 and parental lines. PC2 explains that 20% of the variance separates between parental lines and F4 hybrid.

**Fig. 2.**
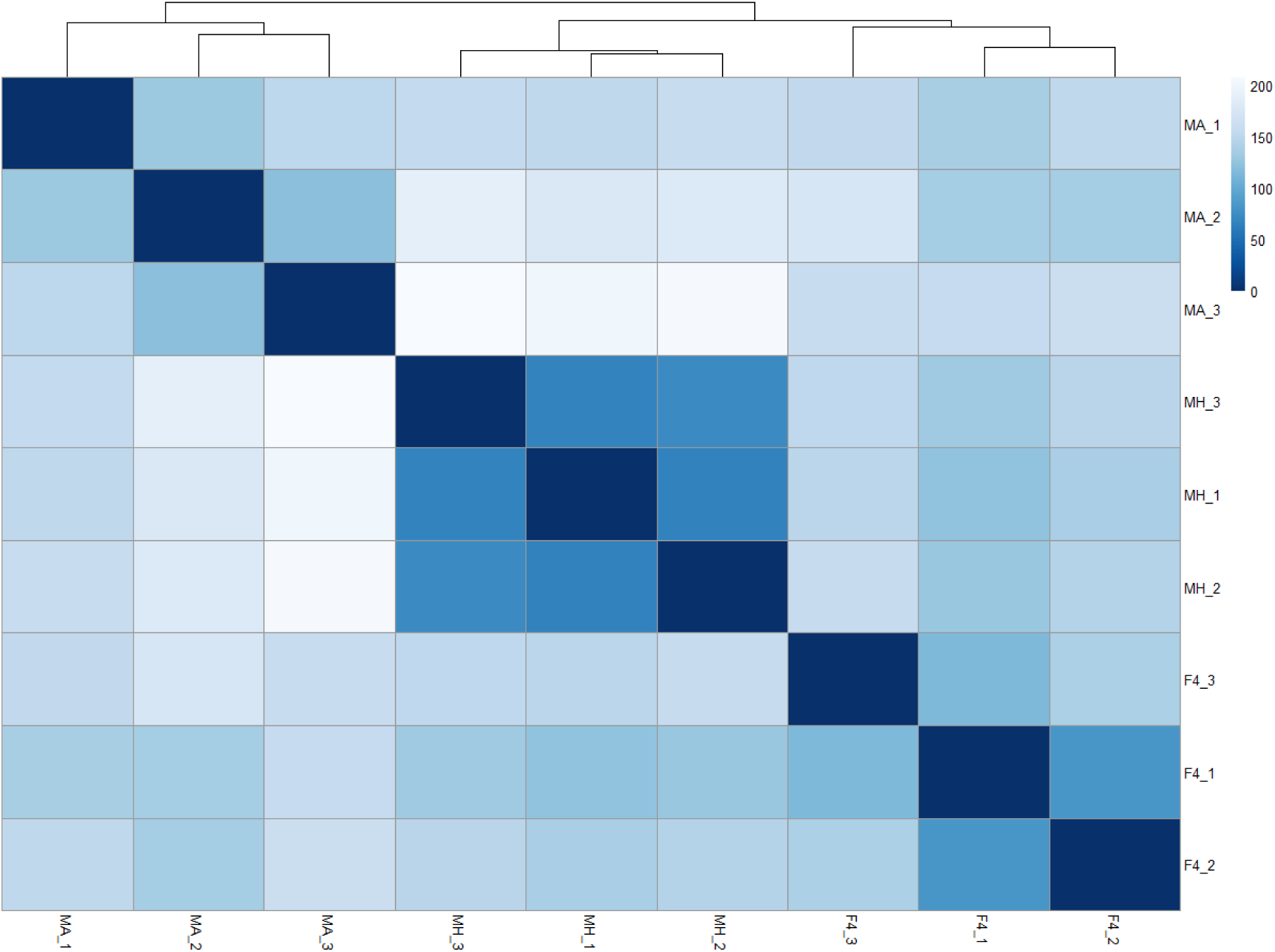
Heatmap of Euclidian distance between samples. Samples (F4 hybrids, Mt. Amasa (MA), and Mt. Hermon (MH) parental lines) from each group were sorted by hierarchical clustering (top). Color corresponds with similarity. Dark blue indicates high similarity, while light blue indicates low similarity.

### Differentially expressed genes between parental lines vs. F4 hybrid

Differential expression analysis was performed using DESeq2. It is important to note that significant differential expression levels between groups were considered to be ∼2-fold or more. In total, 6602 genes showed differential expression between the parental lines and the F4 hybrid. Out of these, 3679 genes showed differential expression between the Mt. Hermon line and the F4 hybrid, while 2311 genes showed differential expression between the Mt. Amasa line and the F4 hybrid. Additionally, 612 genes demonstrated similar expression levels in both parental lines, while different expression levels were observed in the F4 hybrid.

ClusterGap analysis of 6602 differentially expressed genes in parental lines vs. F4 hybrids revealed eight patterns of clusters (Fig. 3). Each cluster represents a group of genes with similar expression patterns in the parental lines and F4 hybrid (Fig. 3). Overall, the eight clusters can be classified into four expression pattern categories: (1) Clusters 1, 3, 4, and 5 consists of 3576 genes that show under-expression in one parental line vs. F4 hybrid (Fig. 3, Fig. 4a). (2) Clusters 2 and 8 consists of 2414 genes overexpressed in one parental line vs. F4 hybrid (Fig. 4, Fig. 5c). (3) Cluster 6 consists of 362 genes under-expressed in both parental lines vs. F4 hybrid (similar expressions in both parents) (Fig. 3, Fig. 4c). (4) Cluster 7 consists of 250 genes overexpressed in both parental lines vs. F4 hybrid (similar expressions in both parents) (Fig. 3, Fig. 4d).

**Fig. 3.**
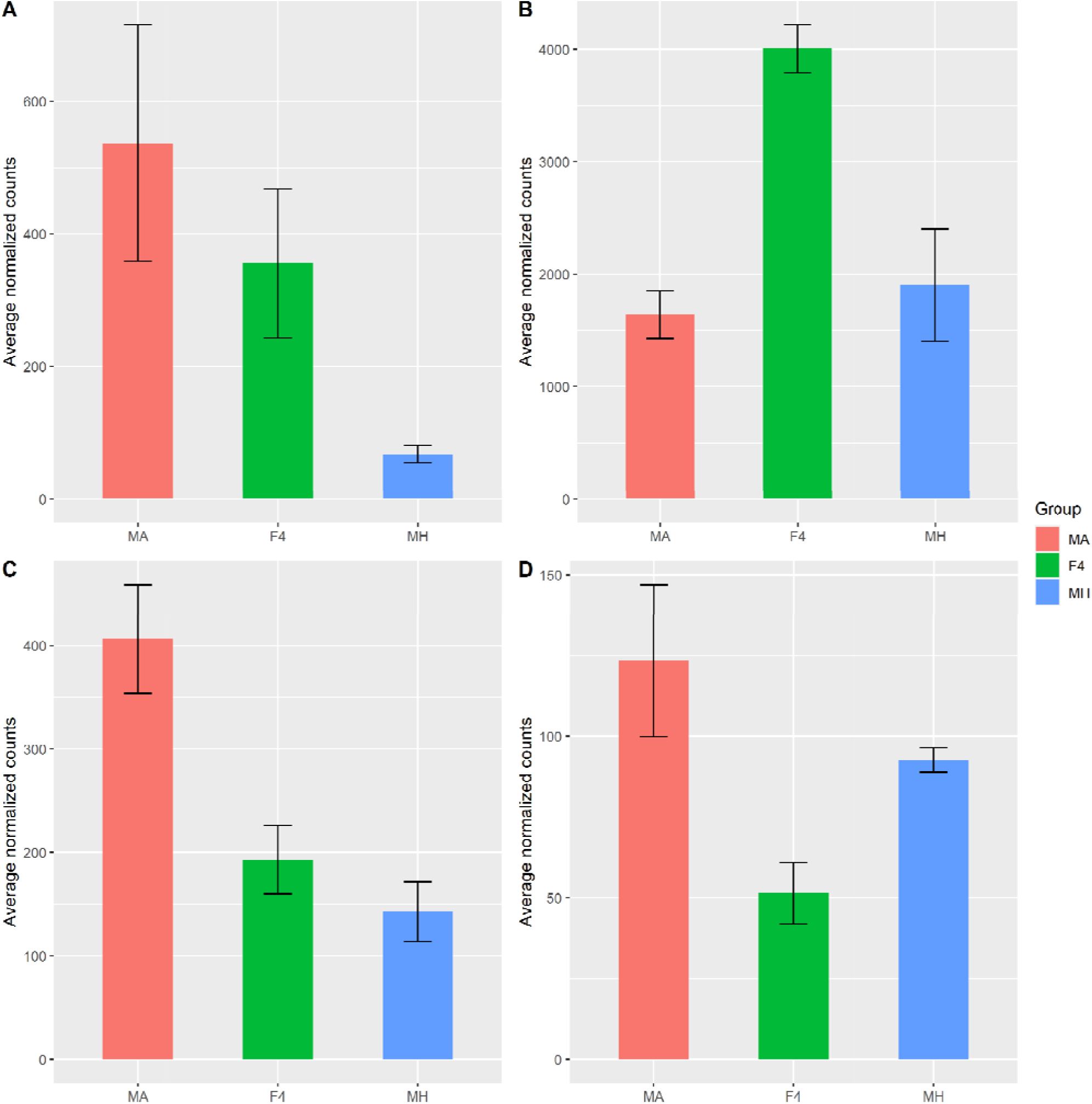
Clustered differentially expressed gene expression by samples. Differential expression analysis revealed a total of 6602 differentially expressed genes among samples. K-mean analysis was performed on a Z-score scaled rlog transformed expression data. Each line represents the mean expression pattern of samples per cluster. Y-axis represents the Z-score scaled rlog transformed expression level. Sample names are on the X-axis. MH – Mt. Hermon samples, MA – Mt. Amasa samples.

**Fig. 4.**
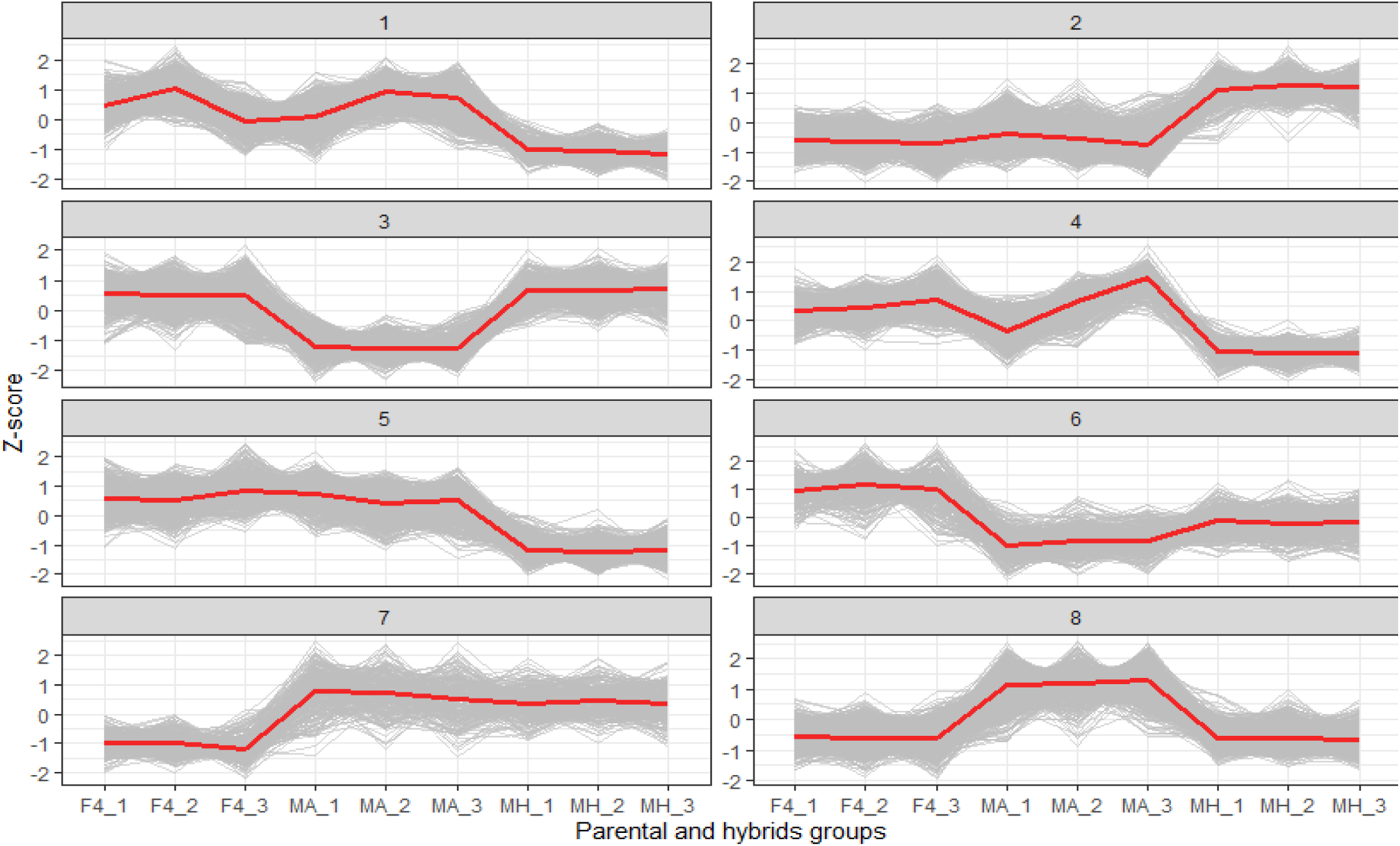
Examples of expression patterns of the four cluster categories. Y-axis notes normalized average counts for each group taken from DESeq2. **A**. Gene TRIDC1AG025480 encodes to a protein associated with plant basic secretory protein (BSP) family related with defense response. This gene belongs to cluster 1, under-expressed genes in Mt. Hermon compared to F4. **B**. Gene TRIDC2BG018130 encodes to a protein associated with photosystem II. This gene belongs to cluster 6, under-expressed in both parental lines compared to F4. **C.** Gene TRIDC1AG036170 is an orthologue of genes encoding to Annexin superfamily protein associated with stress response. This gene belongs to cluster 8, Overexpressed in Mt. Amasa compared to F4. **D.** Gene TRIDC2AG021310 encodes to a protein associated with sucrose synthase activity. This gene belongs to cluster 7, Overexpressed in both parental lines compared to F4.

**Fig. 5.**
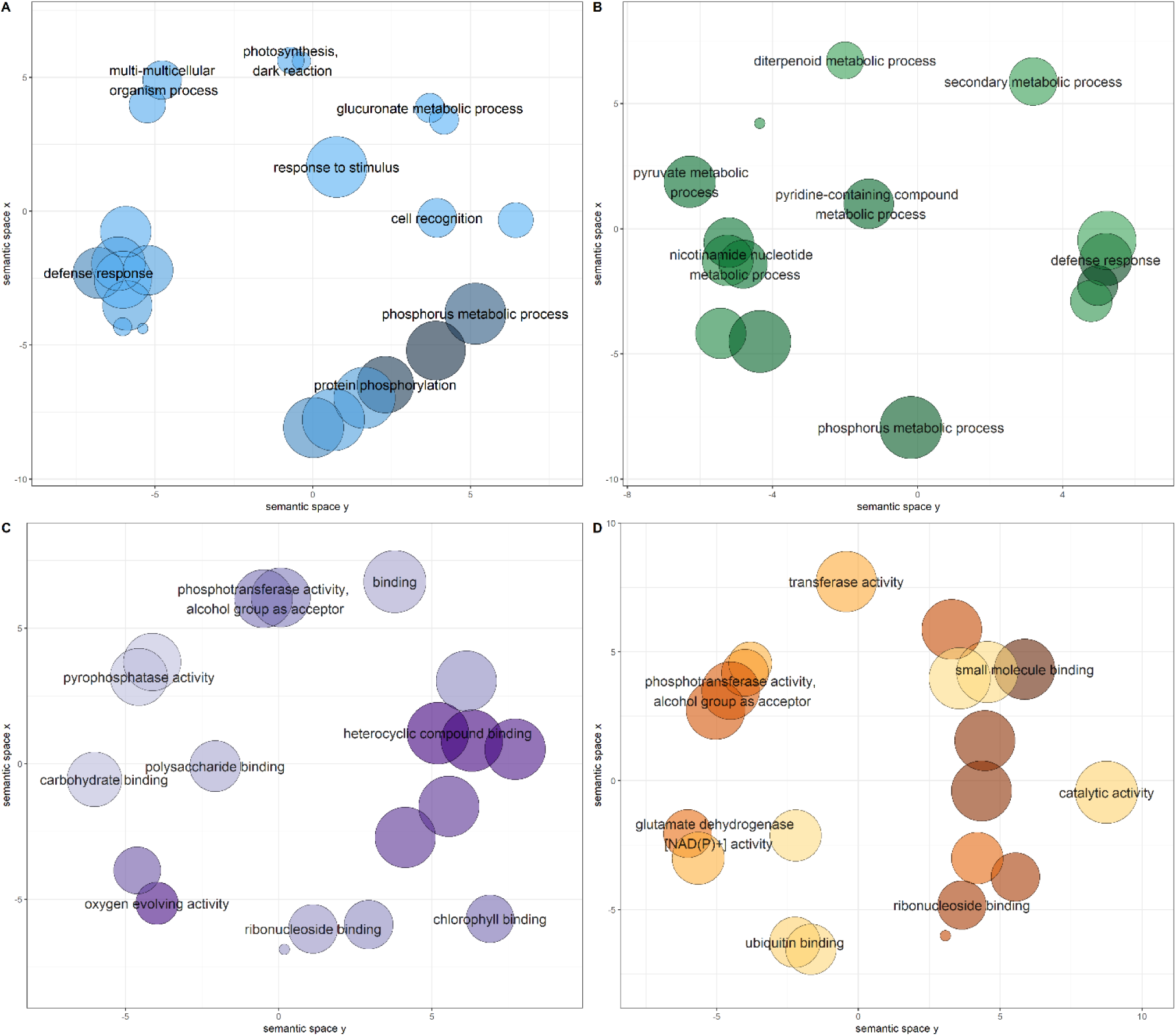
Significantly enriched GO categories were projected onto a two-dimensional semantic space using REVIGO. Color intensity reflects the significance of the enrichment test, with dark colors corresponding to lower P values and white to P closer to 0.05. Circle size indicates the frequency of the GO term in the underlying GOA database (bubbles of more general terms are larger). Revigo assigns names to bubbles representing terms with low dispensability value, meaning non-redundant terms concerning semantically close terms. **A**. BP in cluster 1, Under expressed in Mt. Hermon. **B**. BP in cluster 8, Overexpressed in Mt. Amasa. **C**. MF in cluster 6, Under expressed in both parental lines. **D**. MF in cluster 5, Under expressed in Mt. Hermon.

### SNPs detection between parental lines vs. hybrids

We used the GATK variant calling to identify differences in Single Nucleotide Polymorphisms (SNPs) between the parental lines and their hybrid. After filtering, we obtained 13,170 high-quality SNPs (HQ SNPs) with an overall GQ > 15 across all samples. Out of these, 5,818 SNPs were found in the Mt. Amasa parental line and 7,291 SNPs were found in the Mt. Heromn parental line. Due to the small sample size, we only considered SNPs that matched all three samples per group. In total, we found 1,403 SNPs in 477 differentially expressed genes.

The clusters 1, 4, 5 of Mt. Hermon showed overexpression, while cluster 2 showed underexpression. The largest SNP groups, consisting of 641 and 312 SNPs, were associated with over and under-expressed genes respectively. These SNPs were linked to 273 and 113 genes for over and under-expressed genes, respectively. In Mt. Amasa, clusters 8 and 3 displayed over and under-expressed genes. The SNPs linked with these genes numbered 194 and 151, respectively, and were associated with 73 and 57 genes. F4 had clusters 6 and 7, with upregulated and downregulated genes respectively. These clusters had 38 and 67 SNPs each, linked with 18 genes in each group. SNPeff was used to assign SNPs to their respective genes and then rank each SNP based on its possible effect on those genes. Out of the total SNPs, 642 had a predicted low impact on genes (synonymous SNPs), 303 SNPs had a moderate impact on genes (non-synonymous SNPs), 6 SNPs had a high effect on genes (premature stop codon), and 452 SNPs were predicted to be modifier SNPs (SNPs found in UTR).

After that, we examined whether the SNPs matched the expression pattern revealed by the differential expression analysis. Specifically, SNPs in genes overexpressed in Mt. Hermon (clusters 1, 4, and 5) were only found in Mt. Hermon samples but not in F4 and Mt. Amasa samples, or only in F4 and Mt. Amasa but not in Mt. Hermon. To this end, out of the 1403 SNPs that were observed, 1024 (73%) matched with the expression patterns. Mt. Hermon over and under-expressed clusters had 20-30% unmatched SNPs (clusters 1,4,5, and 2, respectively), while Mt. Amasa over and under-expressed clusters (3 and 8, respectively) had less than 1% unmatched SNPs. The F4 over and under-expressed clusters (6 and 7, respectively) had approximately 90% unmatched SNPs (Fig S2, Additional file 2).

### Gene annotations and functional analysis

We used the gene annotation data, publicly available for WEWSeq 1.0 acc. Zavitan genome, as a reference to analyze the differentially expressed genes in both parental lines and F4 hybrid. To understand the gene functions and biological pathways, we performed a Gene Ontology assessment using SEA analysis in Agrigo v2. To this end, 2680 genes were annotated and significantly classified into; biological processes (BP), cellular components (CC), and molecular function (MF) pathways (Fig. S3, Additional file 2). Overexpressed genes in Mt. Amasa parents (cluster 8) were associated with 25 BP GO terms, while under-expressed genes (cluster 3) were associated with 2 BP, 29 MF, and 7 CC GO terms (Fig. S3, Additional file 2). Overexpressed genes in Mt. Hermon parents (clusters 1,4 and 5) were associated with 152 BP, 73 MF, and 7 CC terms, while under-expressed genes (cluster 2) were associated with 6 BP and 1MF terms (Fig. S3, Additional file 2). Under-expressed genes in the F4 hybrid (cluster 6) were associated with 4 BP, 32MF, and 3 CC terms (Fig. S3, Additional file 2). To visualize the difference in specific GO terms between parental lines and F4 hybrid genotypes, we used over and under-expressed enriched GO categories from clusters 1,2,3,4,5, 6, and 8 to run a REVIGO analysis, which summarizes GO results by removing redundant terms and providing graph-based visualization (21).

In the biological processes class (BP), significant GO terms associated with the under-expressed genes in one parental line vs. F4 hybrid (clusters 1, 3, 4, and 5) were annotated as photosynthesis-related (GO:0019685), dark reaction (6 genes), defense response (GO:0006952, 78 genes), protein phosphorylation (GO:0006468, 194 genes), ncRNA metabolic process (GO:0034660, 89 genes), protein-DNA complex assembly (GO:0065004, 22 genes) and ribonucleoprotein complex biogenesis (GO:0022613, 76 genes) (Fig. 5a, Fig. S4b-d, Additional file 2). Overexpressed genes in one parental line vs. F4 hybrid (clusters 2 and 8) were annotated as response to hexose (GO:0009746), nucleotide phosphorylation (GO:0046939, 15 genes), defense response (GO:0006952, 83 genes), response to hexose (GO:00097460, 13 genes), and defense response to fungus (GO:0050832, 27 genes) (Fig. 5b, Fig. S4a, Additional file 2). Finally, under-expressed genes in the F4 hybrid (cluster 6) were annotated as photosynthesis light reaction (GO:0019684, 15 genes) (Fig. S4e, Additional file 2).

In the molecular function class (MF), significant GO terms associated with under-expressed genes in one parental line vs. F4 hybrid (GO terms were found only in clusters 1, 3, and 5) were annotated as phosphotransferase activity, alcohol group as acceptor (GO: 0016773, 244 genes), purine nucleoside binding (GO:0001883, 481 genes) (Fig. 5d, Fig. S5a,b, Additional file 2). Under-expressed genes in the F4 hybrid (cluster 6) were annotated as electron transporter, transferring electrons within the cyclic electron transport pathway of photosynthesis activity (GO:0045156, 6 genes) and chlorophyll-binding (GO:0016168, 8 genes) (Fig. 4c).

In the cellular component class (CC), GO terms were found only in genes associated with under-expressed genes in one parental line vs. F4 hybrids (GO terms were found only in clusters 1 and 3) and under-expressed genes in F4 hybrid (cluster 6). Genes under-expressed in one parental line were annotated as photosystem I (GO:0009522, 15 genes) and membrane part (GO:0044425, 199 genes), and protein-DNA complex (GO:0032993, 21 genes) (Fig. S6 a,b, Additional file 2). Under-expressed genes in the F4 hybrid were annotated as photosystem II (GO:0009523, 8 genes). Note that a detailed list of all annotated genes can be found in Additional file 3 and Fig. S2. Additional file 2.

### Phenotypic assessments in hybrids vs. parental lines

We analyzed plant height, seed yield, and spike number. The F4 hybrid had intermediate seed yield and spike number but was significantly taller than the parental lines (Fig. S7, Additional file 2). The plant height was measured once a month for three months. Measurements started around Feekes 3.0-4.0 level when at least 3-5 tillers were visible. Height was measured from ground to edge of the highest tiller or edge of the canopy, excluding spikes. Hight measurements of F4 were significantly higher than all the parental lines. In addition, chlorophyll content was measured in fresh leaf tissues. This analysis used nine leaf samples from parental groups and an F4 hybrid (3 biological replicates from each plant). Total chlorophyll content averaged at 0.009±0.006 in Mt. Amasa, 0.1±0.01 in Mt. Hermon, and 0.11±0.01 in F4 hybrid (Fig S8, Additional file 2). To this end, no significant differences were seen between parental lines and the F4 hybrid.

## Discussion

Wheat production has hit breaking record yields thanks to the implementation of modern agricultural and pure lines breeding programs. However, growing worldwide food demand requires constant improvement and increases in wheat production [21]. An important factor hindering the attempt to increase wheat yield is the narrow and homogeneous gene pool among domesticated wheat species, making wheat more susceptible to diseases, pests, and climate change [3–5, 21]. In the past few decades, scientists have looked to the wild ancestors of wheat, like wild emmer wheat, as potential sources for rejuvenating the genetic diversity of cultivated wheat. The significant potential of wild emmer wheat in providing crucial genes for enhanced yield, adaptability to various environmental conditions, and resilience against both biotic and abiotic stressors has gained widespread recognition. In addition, the recent breakthroughs in the whole-genome sequencing [6] and the whole-genome assembly of the wild emmer wheat genome [19] have provided an opportunity to investigate and assess the transcriptomic changes and reveal functional changes within wild emmer wheat.

In this novel research, changes in methylation patterns and transcriptomes were analyzed in two distinct natural populations of wild emmer wheat—Mt. Hermon and Mt. Amasa—as well as in their genetically stable F4 hybrid. These populations represent the cold and hot edge spread of wild emmer wheat populations in Israel and face different environmental challenges.

Understanding the underlying mechanism of these populations’ specific adaptations has been an essential goal in wheat research throughout the years [4, 5, 12, 22]. Our objective was to investigate alterations in methylome and transcriptome levels, as well as identify differentially expressed genes and pathways. This analysis aimed to provide deeper insights into how functional variation contributes to the adaptation of wild emmer wheat to diverse habitats. We observed a notable reduction in methylation levels in the F4 hybrids when compared to the parental lines. Furthermore, we pinpointed 6602 genes that exhibited differential expression between Mt. Hermon, Mt. Amasa, and their F4 hybrid. Gene ontology analysis associated these genes with photosynthesis, defense response, and phosphorylation.

### DNA methylation status in parental lines vs. hybrids

Reduced methylation levels, known as demethylation, have been linked to various factors including responses to environmental stresses, defense mechanisms, activation of transposable elements, and gene expression [23, 24]. The presence of environmental stressors can disrupt the normal growth and development of plants, prompting adjustments in genomic DNA methylation levels as a responsive mechanism to counteract the stress [25, 26]. In rice (Oryza sativa), it has been reported that drought stress triggers widespread alterations in the level of DNA methylation across the genome [26]. In maize, methylation levels were observed to decrease by 1.0–2.2 % under cold stress [25]. A reduction in methylation can also take place following a ‘genomic shock’, as seen in events like hybridization or allopolyploidization [27]. As an illustration, in maize, hybrids derived from two inbred lines exhibited decreased methylation levels in comparison to their corresponding inbred counterparts, with the highest degree of demethylation observed in the hybrids [28]. Similarly, in this study, the methylation levels in the F4 hybrids were found to be lower compared to the parental lines. Upon our evaluation of the overall methylation changes at CCGG sites, we observed a greater number of demethylation events in the F4 hybrid compared to hypermethylation events. Research conducted on soybeans has shown that the greater the reduction in methylation levels, the more pronounced its impact on gene expression levels [29]. In this study, we observed a reduction in methylation levels, leading to a higher occurrence of demethylation events in the F4 hybrids when compared to their parental lines. These results suggest a potential elevation in gene expression. Nevertheless, it’s important to note that we did not directly assess the impact of methylation decrease on gene expression in this study.

### Transcriptome variation

Through a comprehensive examination of genetic diversity and a differential expression analysis, our objective was to discern the influence of each parental lineage on hybrid gene expression. Various models have been proposed to anticipate this effect. The additive genetic model posits that gene expression is equally inherited from both parents, resulting in intermediate expression levels in the hybrid. On the other hand, the dominance genetic model proposes that expression levels are determined by specific alleles, resulting in similarity to one of the parental lineages [30–32]. Additional models, such as the parental effect model, propose that gene expression in hybrids is governed by parental mechanisms like maternal provisioning or genomic imprinting. Consequently, it resembles the gene expression pattern of either the maternal or paternal lineage [31, 33]. Despite our main focus not being the determination of the most fitting model for this case study, it is apparent that the overall genetic diversity of the F4 hybrid lies between that of both parental groups, signifying an intermediate state (Fig. 1). In our examination of differential expression, we found that both the Mt. Amasa group and F4 exhibited a higher count of genes displaying notable expression variations, encompassing both overexpression and underexpression, as opposed to the Mt. Hermon group. Notably, the most prominent cluster of differentially expressed genes was the underexpressed category in Mt. Hermon, which comprised three distinct clusters (1,4 and 5, Fig. 4). This set of 2,433 genes accounts for 37% of the differentially expressed genes. These findings align more closely with the additive or parental effect models rather than the overdominance model. This is because the hybrid exhibited a relatively modest number of differentially expressed genes when compared to both parental lines (clusters 6 and 7). While we may not have definitively identified the most fitting model, it is evident that the F4 hybrid displayed a distinct expression bias towards the paternal line, Mt. Amasa.

### SNPs in differentially expressed genes

Employing SNPs (Single Nucleotide Polymorphisms) is a crucial methodology in genetics, breeding, as well as ecological and evolutionary research. SNPs serve to create genetic markers, uncover cis-regulatory variations, and pinpoint genes linked to particular genetic traits. The utilization of RNA-seq data for SNP identification, although a relatively recent approach, offers several advantages. Firstly, it enables the identification of thousands of SNPs that are correlated with the expression levels of functional genes. Secondly, these SNPs can be linked to biological traits and used to evaluate phenotypes based on genotype. Finally, this method proves to be cost-effective in comparison to other approaches [34, 35].

However, it’s important to note that when utilizing RNA-seq data for variant calling and SNP identification, researchers should be mindful of the potential for false positives and the necessity for high-quality sequencing. The precision of SNP discovery is significantly influenced by factors such as pair-end/single-end sequencing and the length of reads [35].

To mitigate these challenges, we implemented stringent and cautious filtering protocols in this study. Additionally, our analysis was grounded in the well-established relationships between parental groups and hybrids, further bolstering the validity of each identified SNP. Moreover, we consider the identification of SNPs within differentially expressed genes as a preliminary indication, providing a solid foundation for subsequent research and identification efforts. We aimed to determine if SNPs within these genes exhibit a correlation with expression patterns. Our results demonstrated a notable correlation between the expression patterns and the SNPs identified in differentially expressed genes specific to one parental line (clusters 1, 2, 3, 4, 5, and 8). Notably, SNPs in the F4 hybrids consistently originated from one parental line, and we did not observe any entirely new SNPs in the F4 hybrids.

### Hybridization effects

In natural populations, hybridization constitutes a significant evolutionary mechanism capable of augmenting populations. It achieves this by introducing adaptive traits, generating novel lineages, bolstering overall genetic diversity, and engendering heightened phenotypic traits—a phenomenon commonly referred to as heterosis or hybrid vigor [36, 37]. Additionally, hybridization can catalyze the development of more robust reproductive barriers, ultimately culminating in population isolation. This is achieved through the creation of sterile or necrotic hybrids, which hinder gene flow and foster genetic divergence between populations [36, 37]. To assess the functionality of genes impacted by hybridization, we employed Gene Ontology (GO) annotation analysis (Fig. 6, Fig. S3-S6, Additional file 2). Our findings indicate that hybridization between accessions from Mt. Amasa and MH populations may have an impact on defense response, photosynthesis, phosphorylation, as well as DNA and RNA metabolism. Notably, hybrids exhibited expression patterns similar to those of Mt. Amasa in the subset of differentially expressed genes that were under-expressed in Mt. Hermon (clusters 1, 4, 5, Fig. 4). The identified genes were linked to critical processes such as photosynthesis (specifically the dark reaction), defense response, and DNA and RNA metabolism. This suggests that the Mt. Amasa parental line can transmit these beneficial traits through hybridization (Fig 6a, Fig S4c,d, Additional file 2). Conversely, the Mt. Hermon line primarily transmitted traits linked to the MF classification, including purine binding, ATP binding, and ribonuclease binding (Fig. S6b, Additional file 2). In contrast, the F4 hybrid did not exhibit any discernible advantageous traits when comparing differentially expressed genes against both parental lines (cluster 7). Additionally, the F4 hybrid displayed reduced expression levels in genes associated with the photosynthesis light reaction (cluster 6, Fig. S4e), potentially suggesting that the F4 hybrid might not be as proficient as the parental lines.

### Conclusions

To sum up, intraspecific hybridization within natural plant populations, particularly in wild emmer wheat, has been a relatively understudied area. This study delves into the impact of hybridization on methylation alterations, transcriptome-wide changes, differentially expressed genes, and associated pathways. We observed a decrease in methylation levels in hybrids, although we couldn’t directly link this decrease to the observed gene expression patterns. Additionally, we identified notable effects on DNA and RNA metabolism, photosynthesis, and phosphorylation. Notably, most differentially expressed genes related to these traits were predominantly inherited from one parental lineage. Future investigations should focus on understanding how variations in gene expression between hybrids and their parental lines contribute to the adaptation of the nascent hybrid to diverse environmental conditions.

## Methods

### Plant material

Wild emmer wheat accessions from two marginal and natural populations (Mt. Hermon and Mt. Amasa) were chosen for hybridization. These populations represent the most northern population (Mt.Hermon) and the most southern (Mt. Amasa). Previous work in our lab found a population-specific genetic and epigenetic profile in both populations [12]. Accessions from both populations were collected, hybridized, and kindly provided by Dr. Sergei Volis. Reciprocal hybridization was performed by crossing one plant from Hermon and one from Amasa to create the F1 generation. The hybrids underwent selfing to produce subsequent generations, including the F2, F3, and F4 generations. There were two maternal lineages: Mt. Hermon’s maternal lineage and Mt. Amasa’s maternal lineage. Plants were grown in a greenhouse under common garden conditions at Ben-Gurion University between October 2018 to April 2018. All plants throughout all experiments were grown in a 4-liter pot, one for each. The validations of hybrid in the reciprocal crossed were tested by typical AFLP DNA markers (data not shown). Leaf samples for RNA and DNA extraction were collected between the 4^th^ and the 6^th^-week post-germination.

### DNA extraction and Methylation level assessment

Twelve plants were used to assess differences in methylation between parental lines and F4 hybrids, four from three from each of the parental lines, Mt. Hermon and Mt. Amasa, and four from F4 hybrid extraction. DNA was extracted using a DNeasy Plant MINI Kit (Qiagen). DNA quality and concentrations were determined on a 1% gel agarose and by NanoDrop®.

The methylation level was assessed using MSAP (Methylation-Sensitive Amplified Polymorphism). MSAP is a modification of AFLP. It involves two isoschizomers, *HpaII* and *MspI* [12, 38, 39]. Both enzymes cut unmethylated CCGG sites. The genomic restriction fragments are then amplified using two PCR reactions.

*HpaII* and *MspI* differ in their sensitivity to the external or internal cytosine methylation state at the restriction stage. *HpaII* will cleave in cases of external cytosine hemimethylation (only one strand is methylated). *MspI* cleaves when the internal cytosine is methylated. MSAP creates a pattern of monomorphic bands between the *HpaII* and *MspI* isoschizomers. If both isoschizomers digest the DNA templates (from the same DNA sample), it indicates that the CCGG site is unmethylated, while polymorphic bands indicate methylated sites. After the restriction stage, fragments were ligated to adaptor DNA, which includes an overhang complementary to the overhang produced by the restriction enzyme samples. Samples were then amplified twice: The first PCR amplification was a non-selective amplification using primers complementary to the adaptor sequence. The second PCR amplification was a selective amplification using the same primers from the non-selective amplification but with the addition of three random nucleotides. The methylation level of an individual was measured as the number of polymorphic bands between the *MspI* and *HpaII* MSAP reactions in the same individual out of the total number of MSAP bands. Primer sequences used in this study are shown in Additional File 1.

Selective amplification products were electrophoresed in a 3730xl DNA analyzer (Applied Biosystems) and analyzed using GeneMapper v6.0 (Applied Biosystems). Results from GeneMapper were transferred to an Excel table that summarized the presence (1) / absence (0) of bands at every site in all samples. The methylation level was calculated manually as the number of polymorphic bands between *HpaII* and *MspI* patterns divided by the total number of bands (loci) in the two patterns.

### RNA extraction and sequencing

Overall, nine plant samples were selected for RNA-Seq analysis; three from each one of the parental lines, Mt. Hermon and Mt. Amasa, and three from the F4 hybrid. RNA was extracted using a ZR Plant RNA mini-prep kit (Zymo Research, Irvine, USA). Samples were sent to the Technion genome center (Haifa, Israel) for sequencing. RNA quality was assessed using Agilent Bioanalyzer. RNA Libraries were prepared using Illumina TruSeq RNA Library Preparation Kit v2. (Illumina). RNA sequencing was performed on two lanes on the Illumine Hiseq2500 machine at a read length of 50 bp SR. Quality control was evaluated using FASTQC v0.11.5 [40]. RNA-seq reads were aligned against wild emmer wheat reference genome WEWSeq 1.0 acc. *Zavitan* [19] using Salmon [41]. Tximport [42] was used to aggregate transcript-level counts and abundances into gene-level counts and abundances before normalization and differential expression analysis.

### Differential gene expression analysis

Normalization and differential gene expression analysis were conducted using the ‘DESeq2’ R package [43]. Differential gene expression pairwise comparison (e.g., F4 Vs. MH group) was performed using a Wald test (adjust p-value > 0.05). Genes with fold-change > 2 were considered viable for further analysis. Differentially expressed gene counts were transformed using rlog transformation. Rlog transformation transforms the normalized counts to the log2 scale to minimize differences between samples with low and high counts.

### SNPs and variant calling

Variant calling was performed to observe if there is a possible effect of Single Nucleotide Polymorphism (SNP) on the observed DEGs. Cleaned reads were mapped using the STAR program [44]. SNPs were identified using the Genome Analysis Toolkit (GATK, version 4.0.5.0) [45] recommended pipeline, with hard SNPs filtering as an exception. SNP variant calling was done using HaplotypeCaller and GenotypeGVCFs packages of GATK.

As wild emmer wheat is not liable for the GATKs VQSR filter tool, high-strength filtering based on GATK-recommended parameters was done. Each parameter threshold was determined following data distribution and used stricter thresholds than GATK-recommended thresholds. Credible variants were defined as variants that satisfied the following parameters: quality depth (QD) <2.0, FisherStrand (FS)>60.0, RMSMappingQuality (MQ)<40.0, and ReadPosRankSum < −8.0.

Filtered SNPs were annotated using SNPeff [46]. SNPeff assigns SNPs to their respective gene and predicts the possible effect onset genes (e.g., change in amino acids) [46]. SNPs were then uploaded to R for further analysis. Additional filtration was done using the GQ (Genotype quality) parameter. SNPs with GQ higher than 15 across all samples were used for further analysis. SNP’s homo/heterozygosity for each sample was determined using AB (allelic balance) parameter. SNPs with AB between 0.2 to 0.8 were considered heterozygotes. Finally, we examined any correlation between SNPs’ appearance in differentially expressed genes and the expression level in each group (parental and hybrids). For example, SNPs in genes overexpressed in one parental group could only be found in that set group but not in hybrids or the other parental group or vice versa, these SNPs could only be found in hybrids and the other parental group.

### Gene functional classification and pathway analysis

Genes were annotated using matched orthologue genes using *Ensemblplants* Biomart [47] and BLASTn (E-value cutoff of 10^-10^, Additional file 3) against bread wheat (*Triticum aestivum*), Arabidopsis (*Arabidopsis thaliana*), and rice (*Oryza sativa Japonica*). AgriGO v2 [48] was used to perform Gene Ontology (GO) singular enrichment analysis (SEA) for all differentially expressed genes. GO is a functional tool that classifies and characterizes information on an attribute of gene products in distinct, not overlapping, groups [48]. GO analysis takes a group of genes associated with a specific criterion (e.g., similar gene expression). It identifies biological processes, cellular components, or molecular functions overrepresented in that group compared to a background population from which the query list is derived. Differentially expressed genes (DEGs) were used as gene groups while the wild emmer wheat reference genome WEWSeq 1.0 acc. *Zavitan* [19] was used as the background population. Reduction and Visualization GO analyses were performed using REVIGO (http://revigo.irb.hr/). REVIGO removes redundant GO terms using SimRel semantic similarity measure (the analysis allowed for a medium level of similarity, set at 0.7.) [49].

### Statistical analysis

Pairwise comparison between two specific groups (for example, F4 to MH group) was performed using a Wald test (adjust p-value > 0.05).

To observe the overall genetic distance between and within sample groups, we used Principal component analysis (PCA) and heatmap.

PCA observed the overall genetic distance between samples. It is a technique used to highlight variation in data and flash out strong patterns in multi-dimensional data sets such as transcriptomic data Fields[50]. The output of a PCA transformation can be used to make data easy to explore and visualize. PCA projects data onto a two-dimensional (or three-dimensional in some cases) plane so that they spread out in the two directions that explain most of the variance between samples [51]. The first principal component (PC1) explains the most significant possible variance in the data (e.g., 60% of the variance between control and tested samples is explained by the genes defining PC1). PC2 explains the second most significant portion of variance, and so on. Traditionally, the first few PCs are used for visualization since they capture most variation from the original data set [52].

PCA plot was conducted using log-transformed data. PCA graph was plotted using the ggplot2 package in R [53].

Transformed DEGs counts were scaled using Z-score (standard score) To perform K-mean cluster analysis and heatmaps of differentially expressed genes. Scaling was performed to identify clusters of genes with similar expression profiles rather than similar expression levels. A total samples expression level heatmap was constructed using the ‘pheatmap’ package in R [54]. Samples were sorted by hierarchical clustering using the default clustering provided by the pheatmap package.

DEGs were clustered according to their expression level across all samples using K-mean. The k-mean cluster uses an a priori number of clusters to split the data accordingly. The number of clusters (the number of Ks) was determined by applying Gap statistics [55] using the clusGap function in R [56]. K-means clustering was performed according to the number cluster using the K-means base function in R. Clusters were then subjected to gene ontology analysis using AgriGO.

## Declarations

## Ethics approval and consent to participate

Not applicable

## Consent for publication

Not applicable

## Availability of data and materials

All data generated or analyzed during this study are included in this published article [and its supplementary information files]

## Authors’ contributions

A.Z: generated data, data analysis, wrote manuscript. K.K: corresponding author, prepared and submitted manuscript. All authors contributed to the article and approved the submitted version

## Acknowledgments

We would like to thank Vadim Khasdan for his assistance with the manuscript preparation. This work was supported by a grant from the Israel Science Foundation (grant# 1311/21) to K. K.

## Abbreviations

AB: Allelic balance
BP: Biological processes
CC: Cellular components
DEGs: Differentially expressed genes
GO: Gene ontology
GQ: Genotype quality
IPCC: Intergovernmental Panel on Climate Change
MA: Mt. Amasa
MF: Molecular function
MH: Mt. Hermon
MSAP: Methylation-Sensitive Amplified Polymorphism
PC: Principal component
PCA: Principal component analysis
SNP: Single Nucleotide Polymorphism

