## Supplementary material for "Transcriptome variations in hybrids of wild emmer wheat (*Triticum turgidum ssp. dicoccoides*)": tables s1 and s2-additional file 1

Supplementary Table S1. Adaptors and primers used in MSAP reaction

| Eco-adapter1 | 5' CTC GTA GAC TGC GTA CC 3’ |
| --- | --- |
| Eco-adapter2 | 5’ AAT TGG TAC GCA GTC TAC 3’ |
| Eco-pre | 5' GAC TGC GTA CCA ATT CA 3’ |
| E-ACT | 5' GAC TGC GTA CCA ATT CAC T 3' |
| E-ACA | 5' GAC TGC GTA CCA ATT CAC A 3' |
| E-ACC | 5' GAC TGC GTA CCA ATT CAC C 3' |
| H/M- adapter 1 | 5' GAT CAT GAG TCC TGC T 3' |
| H/M- adapter 2 | 5' CGA GCA GGA CTC ATG A 3' |
| H/M-pre | 5' ATC ATG AGT CCT GCT CGG 3' |
| H/M-TCAA | 5' CAT GAG TCC TGC TCG GTC AA 3' |

Supplementary Table S2**.** Cytosine methylation levels in parental lineages and hybrids

| **Primer**  **combination** | **Parental/hybrid group** | **MH** | **MA** | **F4** |
| --- | --- | --- | --- | --- |
| E-ACC | Average No. of Bands | 118.2 | 156.3 | 161.3 |
|  | Average polymorphic bands | 102.2 | 132.3 | 130.8 |
|  | Average Polymorphism | 86.3 | 84.6 | 81.2 |
|  | standard error | 1.7 | 1.0 | 3.0 |
| E-ACA | Average No. of Bands | 100.8 | 131.7 | 133.7 |
|  | Average polymorphic bands | 80.8 | 98.0 | 99.3 |
|  | Average Polymorphism | 80.3 | 74.4 | 75.0 |
|  | standard error | 1.7 | 2.4 | 1.3 |
| E-ACT | Average No. of Bands | 79.0 | 81.8 | 87.3 |
|  | Average polymorphic bands | 71.4 | 71.8 | 62.0 |
|  | Average Polymorphism | 90.3 | 88.1 | 71.2 |
|  | standard error | 2.1 | 3.3 | 2.2 |
| Average | **Average No. of Bands** | **99.3** | **123.2** | **127.4** |
|  | **Average polymorphic bands** | **84.8** | **100.7** | **97.4** |
|  | **Average Polymorphism** | **85.6** ^α^ | **82.4** ^α^ | **75.8** ^β^ |
|  | **standard error** | **1.8** | **2.2** | **2.2** |

α and β note statistical significance
