## Supplementary material for "Transcriptome variations in hybrids of wild emmer wheat (*Triticum turgidum ssp. dicoccoides*)": figures s1-s6-additional file 2

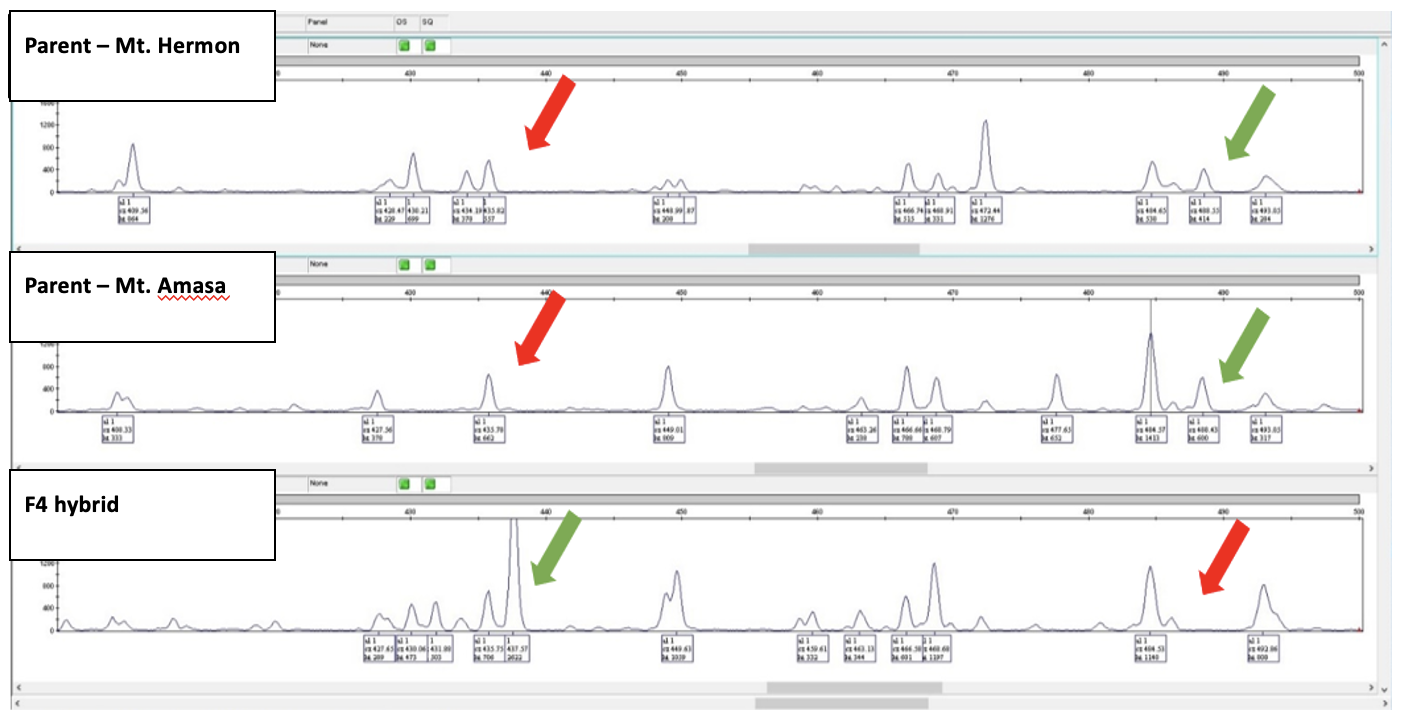

**Fig. S1.** An example of novel and absent peaks (bands) among Hermon maternal lineage offspring. Parental plants (Hermon and Amasa) and offspring are hybrid from generation F3. The green arrows indicate the presence of methylation sites, and the red arrows indicate the absence of methylation sites

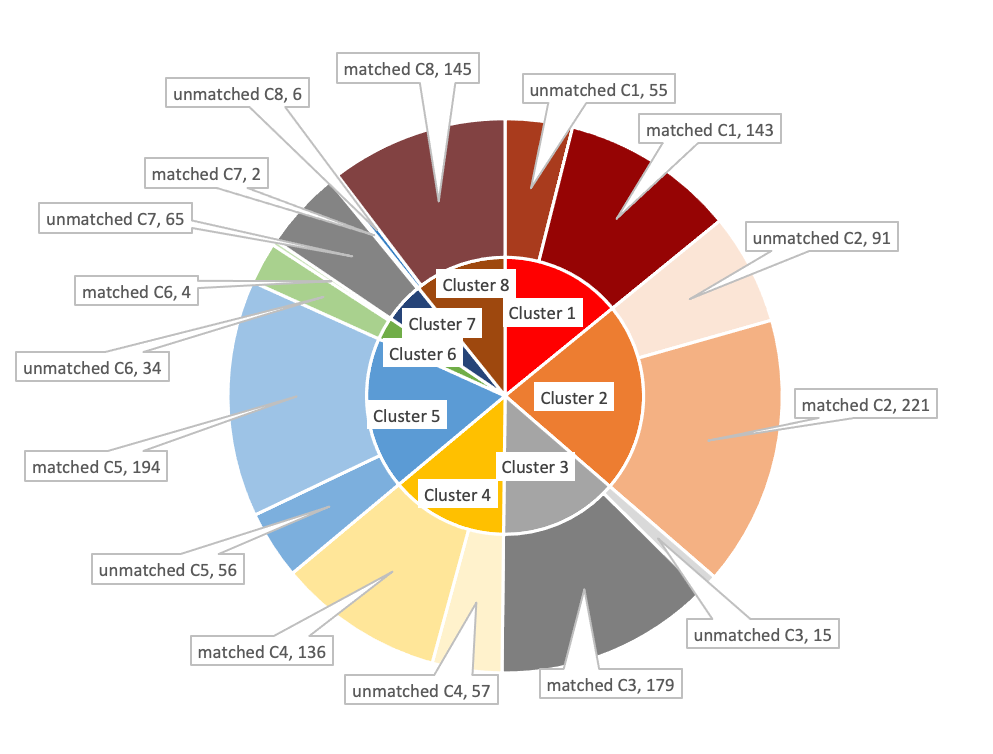

**Fig. S2***.* SNPs divided according to DE analysis, matched SNPs fit the expression pattern of their cluster, while unmatched SNPs do not fit their expression patterns. MH clusters (overexpressed genes: 1,4,5 and under-expressed genes:2) had between 20-30 % unmatched SNPs, while MA clusters (over-expressed: 3 and under-expressed 8) had less than 1% unmatched SNPs. F4 clusters (6,7) had approximately 90% unmatched

|  |  |  |
| --- | --- | --- |
| Cluster 1 | 198 |  |
| Cluster 2 | 312 |  |
| Cluster 3 | 194 |  |
| Cluster 4 | 193 |  |
| Cluster 5 | 250 |  |
| Cluster 6 | 38 |  |
| Cluster 7 | 67 |  |
| Cluster 8 | 151 |  |
| unmatched C1 |  | 55 |
| matched C1 |  | 143 |
| unmatched C2 |  | 91 |
| matched C2 |  | 221 |
| unmatched C3 |  | 15 |
| matched C3 |  | 179 |
| unmatched C4 |  | 57 |
| matched C4 |  | 136 |
| unmatched C5 |  | 56 |
| matched C5 |  | 194 |
| unmatched C6 |  | 34 |
| matched C6 |  | 4 |
| unmatched C7 |  | 65 |
| matched C7 |  | 2 |
| unmatched C8 |  | 6 |
| matched C8 |  | 145 |

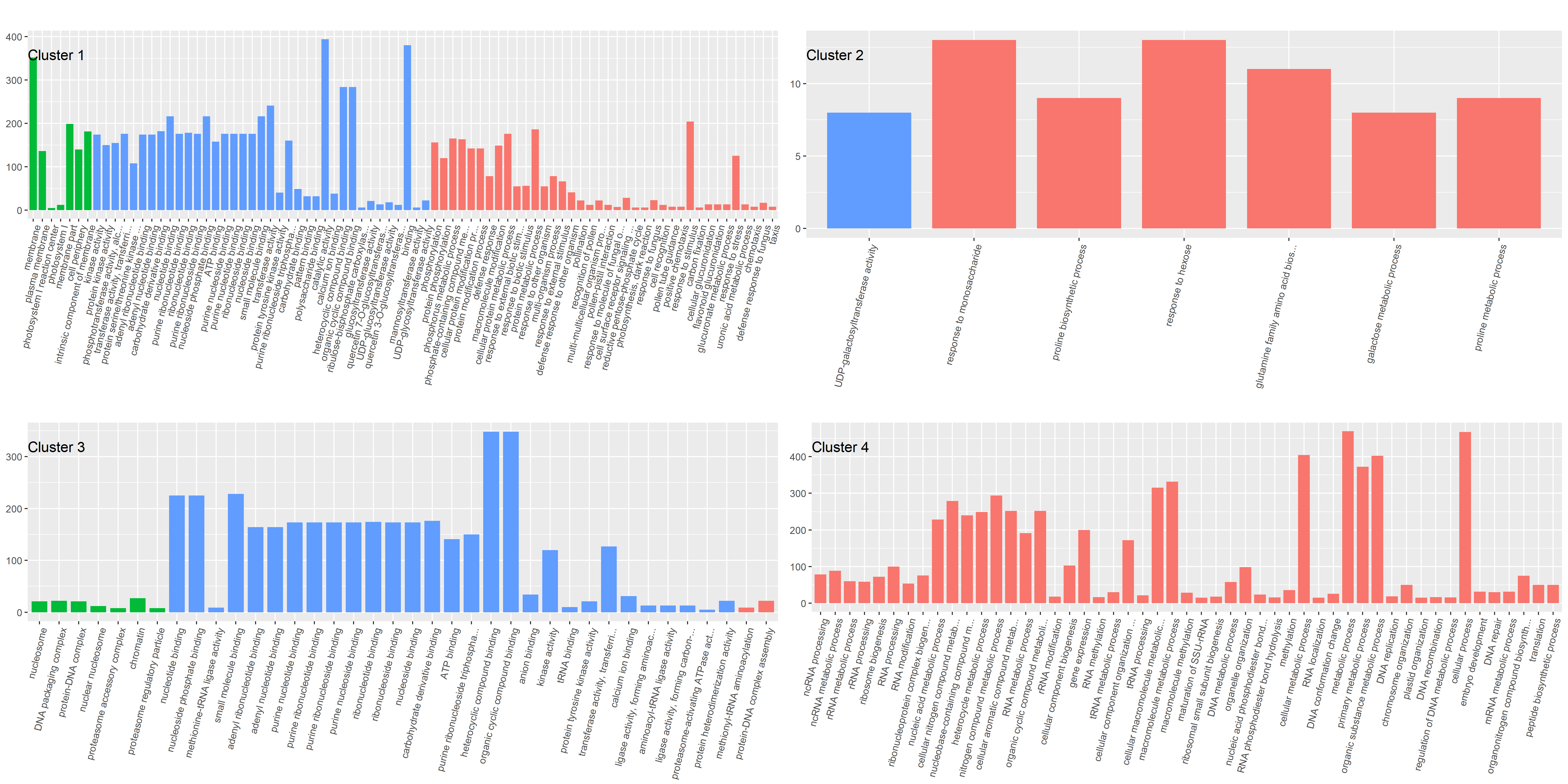

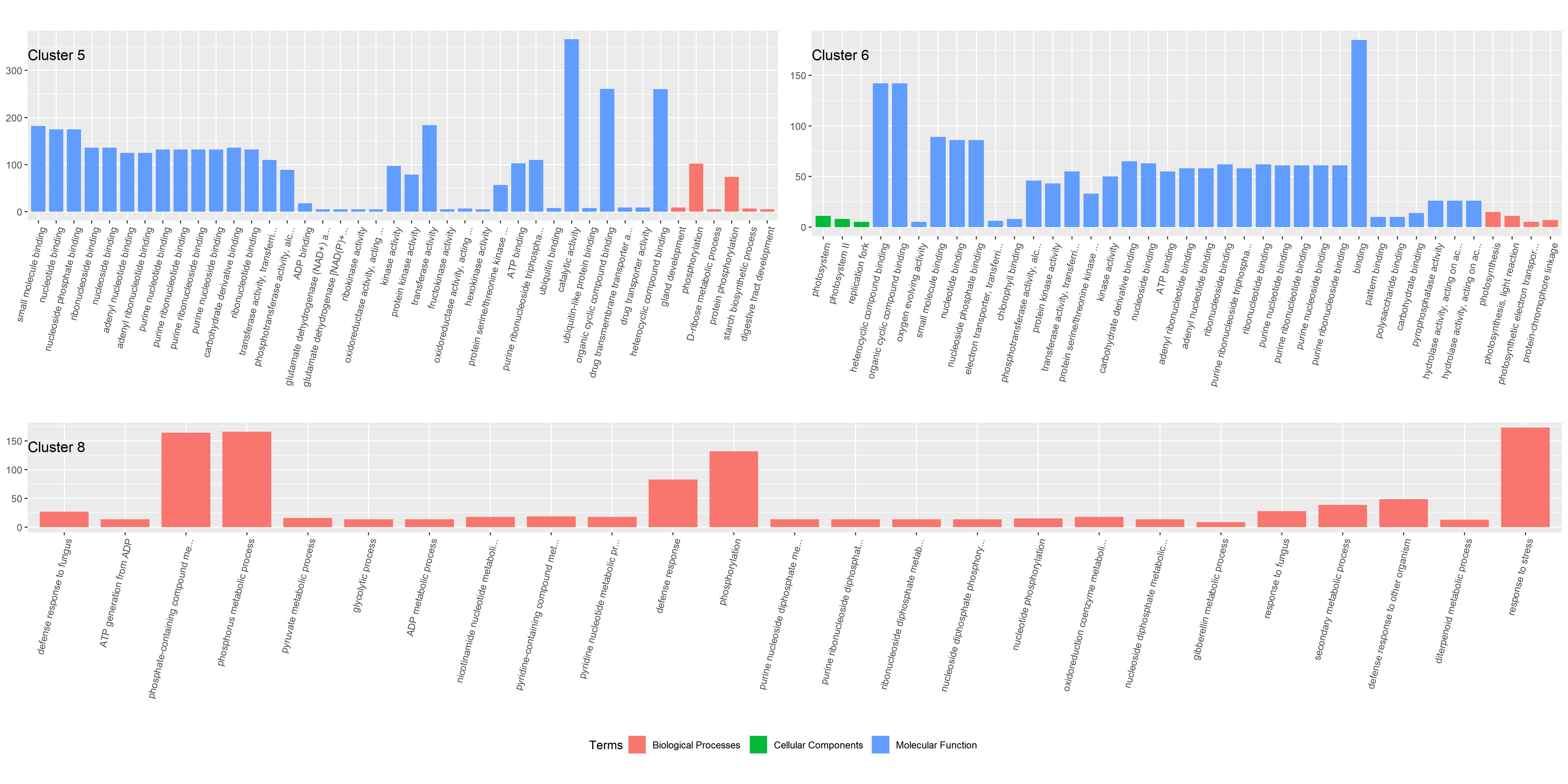

**Fig. S3.** Gene ontology (GO) enrichment analysis diagram for biological processes (BP), cellular components (CC), and molecular function (MF) in clusters 1,2,3,4,5,6,8. Note that cluster 7 did not yield statistically significant GO terms. The X axis represents the number of genes in represented by each term.

**Fig. S4.** Significantly enriched biological process (BP) GO categories were projected onto a two-dimensional semantic space using REVIGO. Color intensity reflects the significance of the enrichment test, with dark colors corresponding to lower P values and white to P closer to 0.05. Circle size indicates the frequency of the GO term in the underlying GOA database (bubbles of more general terms are larger). Revigo assigns names to bubbles representing terms with low dispensability value, meaning non-redundant terms with respect to semantically close terms. **A.** BP in cluster 2, Over express in Mt. Hermon. **B.** BP in cluster 3, Under expressed in Mt. Amasa. **C.** BP in cluster 4, Under expressed in Mt.Hermon. **D.** BP in cluster 5, Under expressed in Mt.Hermon. **E.** BP in cluster 6, Under expressed in both parental lines

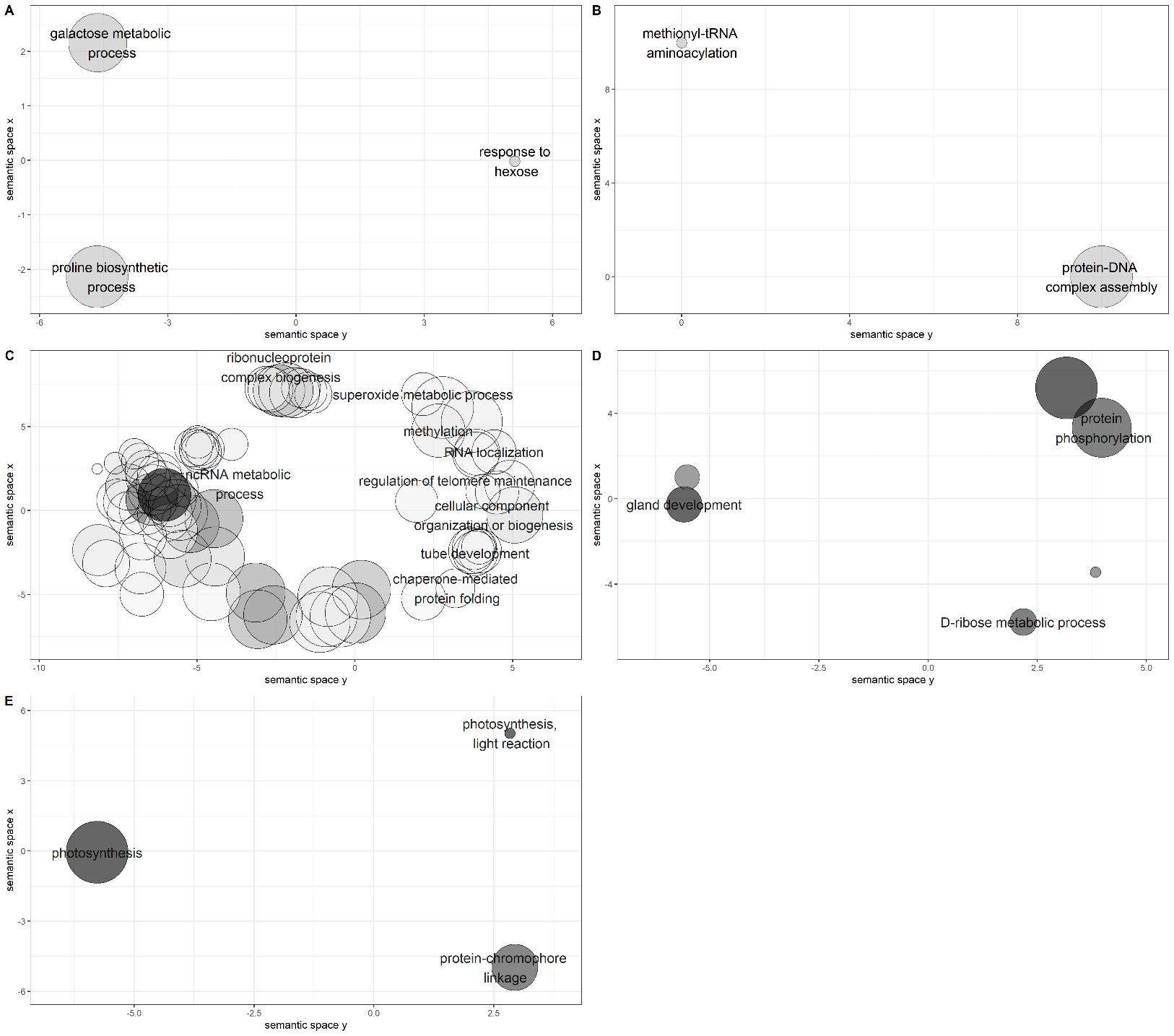

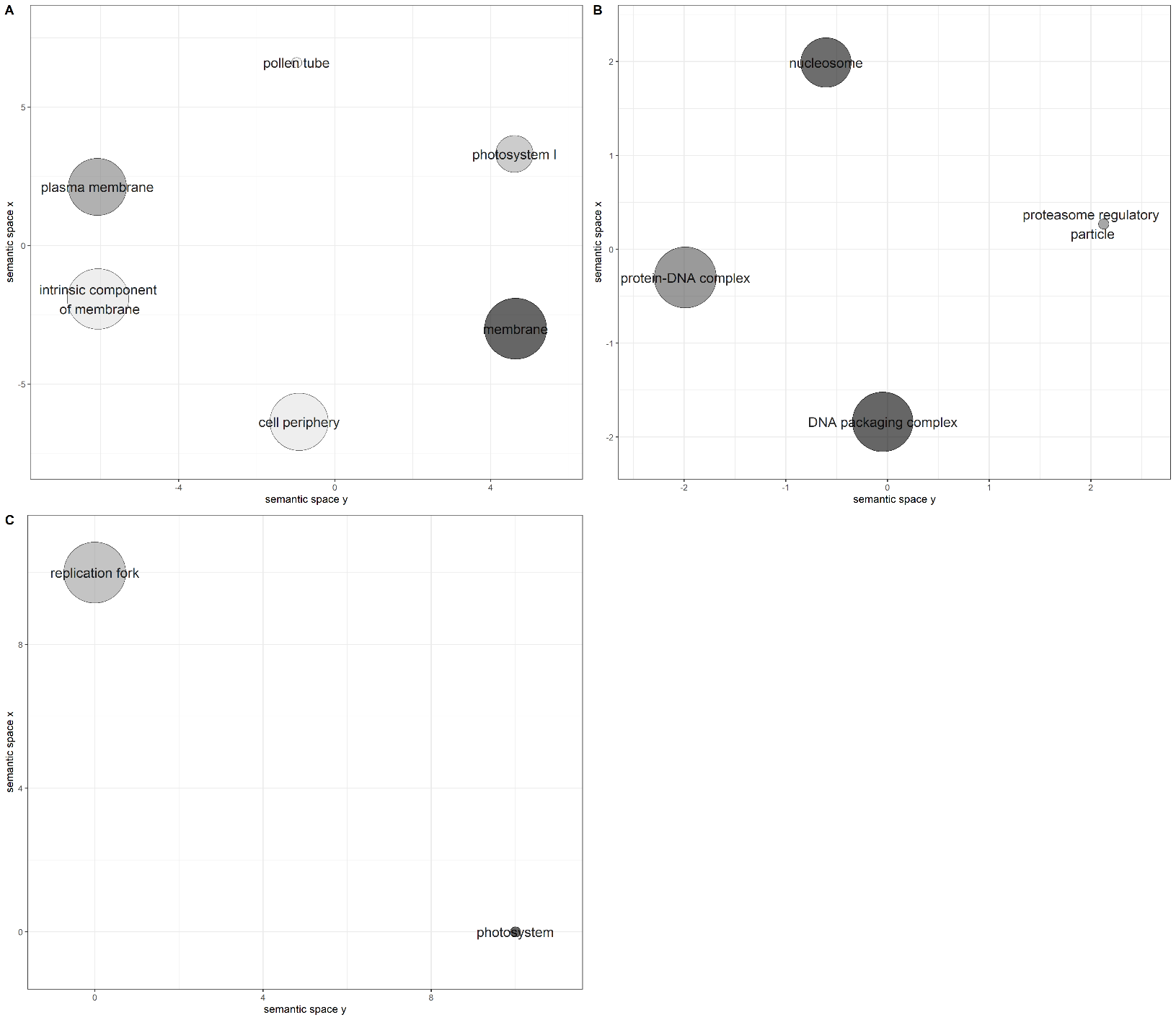

**Fig. S5.** Significantly enriched cellular components (CC) GO categories were projected onto a two-dimensional semantic space using REVIGO. Color intensity reflects the significance of the enrichment test, with dark colors corresponding to lower P values and white to P closer to 0.05. Circle size indicates the frequency of the GO term in the underlying GOA database (bubbles of more general terms are larger). Revigo assigns names to bubbles representing terms with low dispensability value, meaning non-redundant terms with respect to semantically close terms. **A.** CC in cluster 1, Under expressed in Mt.Hermon. **B.** CC in cluster 3, Under expressed in Mt. Amasa. **C.** CC in cluster 6, Under expressed in both parental lines.

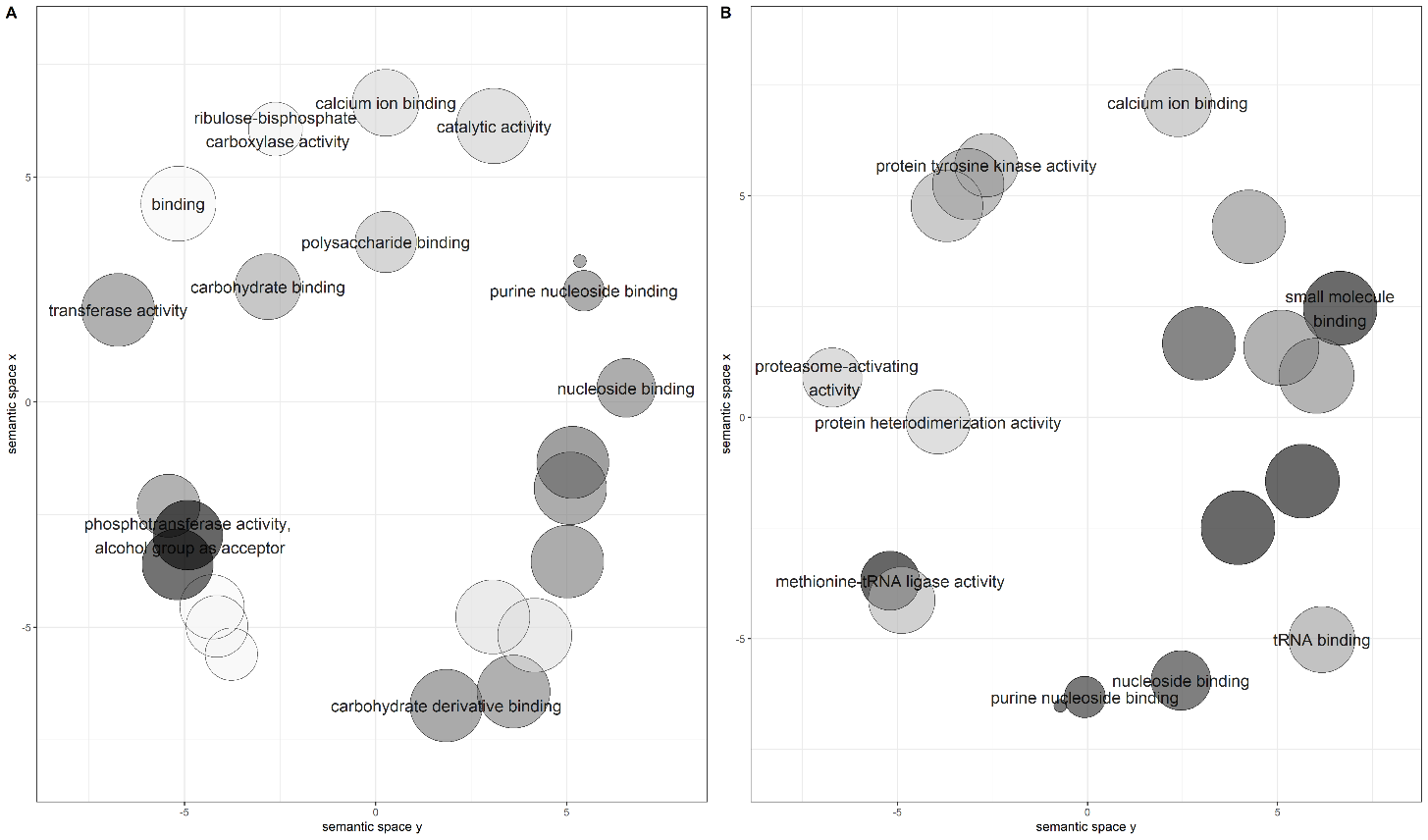

**Fig. S6.** Significantly enriched Molecular function (MF) GO categories were projected onto a two-dimensional semantic space using REVIGO. Color intensity reflects the significance of the enrichment test, with dark colors corresponding to lower P values and white to P closer to 0.05. Circle size indicates the frequency of the GO term in the underlying GOA database (bubbles of more general terms are larger). Revigo assigns names to bubbles representing terms with low dispensability value, meaning non-redundant terms with respect to semantically close terms. **A.** MF in cluster 1, Under expressed in Mt.Hermon. **B.** MF in cluster 3, Under expressed in Mt. Amasa.

.

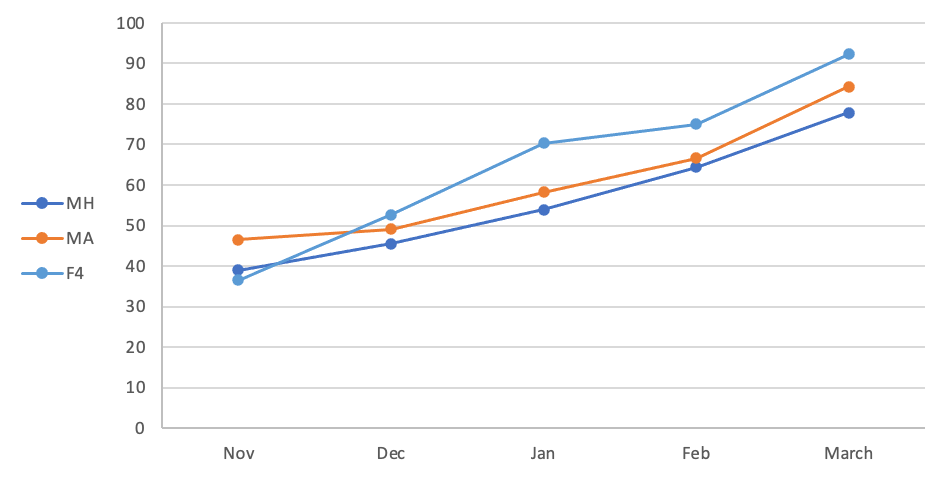

**Fig. S7.** Height measurements of parental groups and hybrids. Measurements started around Feekes 3.0-4.0 level when at least 3-5 tillers were visible. Height was measured from ground to edge of the highest tiller or edge of the canopy, excluding spikes.

|  | MH | MA | F4 |
| --- | --- | --- | --- |
| Nov | 38.9667 | 46.5 | 36.5 |
| Dec | 45.4643 | 49.0455 | 52.7 |
| Jan | 53.8571 | 58.1818 | 70.25 |
| Feb | 64.3214 | 66.6 | 75 |
| March | 77.875 | 84.2 | 92.25 |

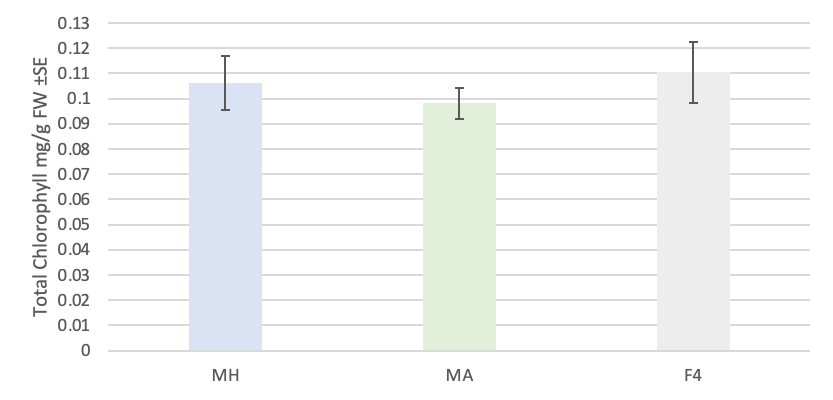

**Fig. S8.** Total chlorophyll content in leaves of parental groups and F4 hybrid (mean ± SE, n = 9).
No significant differences were found between all three groups

|  | Total Chlorophyll mg/g FW ±SE |
| --- | --- |
| MH | 0.106087 |
| MA | 0.0981072 |
| F4 | 0.110348 |
